# Greater breadth of vaccine-induced immunity in females than males is mediated by increased antibody diversity in germinal center B cells

**DOI:** 10.1101/2022.05.31.494237

**Authors:** Rebecca L. Ursin, Santosh Dhakal, Hsuan Liu, Sahana Jayaraman, Han-Sol Park, Harrison R. Powell, Morgan L. Sherer, Kirsten E. Littlefield, Ashley L. Fink, Zexu Ma, Alice L. Mueller, Allison P. Chen, Yishak A. Woldetsadik, Patricia J. Gearhart, H. Benjamin Larman, Robert W. Maul, Andrew Pekosz, Sabra L. Klein

## Abstract

Inactivated influenza vaccines induce greater antibody responses in females than males among both humans and mice. To test the breadth of protection, we used recombinant mouse-adapted A/California/2009 (maA/Cal/09) H1N1 viruses containing mutations at one (1M), two (2M), or three (3M) antigenic sites, in addition to a virus containing the 1M mutation and a substitution of the Ca2 antigenic site (Sub) with one derived from an H5 hemagglutinin (HA) to challenge mice of both sexes. Following maA/Cal/09 vaccination, females produced greater virus-specific class-switched IgG and IgG2c antibodies against the vaccine and all mutant viruses, and antibodies from females recognized more unique, linear HA epitopes than antibodies from males. While females had greater neutralizing antibody (nAb) titers against the vaccine virus, both sexes showed lower neutralization capacity against mutant viruses. After virus challenge, vaccinated females had lower pulmonary virus titers and reduced morbidity than males against the 1M and 2M viruses, but not the Sub virus. Females generated greater numbers of germinal center (GC) B cells containing superior somatic hypermutation frequencies than vaccinated males. Deletion of activation-induced cytidine deaminase (*Aicda)* eliminated female-biased immunity and protection against the 2M virus. Harnessing methods to improve GC B cell responses and frequencies of somatic hypermutations, especially in males, should be considered in the development of universal influenza vaccines.

**Summary:** Compared with males, inactivated influenza vaccination of female mice causes greater production of class-switched, somatically-hypermutated antibodies and a more robust germinal center B cell response, leading to more diverse H1N1 antigen recognition and better protection against mutant influenza A viruses.

## Introduction

Influenza virus infection is a major public health threat, and prior to the COVID-19 pandemic, was responsible for over 5 million cases and around a quarter of a million deaths worldwide each year (1, 2). Infection with influenza A (IAV) and B viruses is seasonal and the pathogenesis, antigenicity, and virulence of these viruses can change each season. Influenza viruses have segmented, RNA genomes which enable the viruses to become antigenically distinct and evade immunological memory (2-4). The propensity for influenza viruses to undergo antigenic drift results in the need for annual updates to the seasonal influenza vaccine virus strain selection.

Annual vaccination prevents severe disease and the spread of the virus; the available vaccination platforms, however, can vary widely in their efficacy and protection (5, 6). Most of the worldwide population receives an inactivated influenza vaccine (IIV) that targets the influenza attachment protein, hemagglutinin (HA). IIVs include either three (trivalent: one H1N1, one H3N2, and one influenza B) or four (quadrivalent: the same as trivalent but with both influenza B lineages) strains of influenza, but each strain can independently vary in efficacy and induction of immunological protection (7, 8). For example, during the 2017-18 influenza season, a mismatch in the H3N2 circulating and selected vaccine strains was associated with mutations introduced by egg adaptation of the vaccine virus, which led to an influenza season that was as severe as the 2009 H1N1 pandemic season in the United States (9).

Improved influenza vaccine efficacy requires a better understanding of both viral and host factors that affect vaccine-induced immune responses (10-12). Viral factors include mutations introduced into the vaccine strains during egg adaptation and vaccine manufacturing, seasonal antigenic drift and shift, immunological pressure from the host, and post-translational modifications (e.g. addition or loss of glycosylation sites) of the virus (7, 13). There are many biological host variables that influence vaccine efficacy and protection, including but not limited to biological sex, age, reproductive status, and body mass index (14, 15). Biological sex, in particular, is an important predictor of antibody responses and protection following receipt of influenza vaccines, with females generally producing greater antibody responses than males. These observed sex differences are age dependent and associated with circulating sex steroid hormone concentrations (16, 17).

For many vaccines, including IIV, the primary correlate of protection is the antibody response generated by B cells (6). After immunization with IIV, female mice exhibit greater quantity and quality of vaccine-specific antibodies and protection from both infection and severe disease than males (9, 18-21). While being female is a predictor of greater immunity and protection against influenza, few studies have considered sex differences in protection against influenza in the context of evolved virus mutations, which we have shown is a limitation to female-biased protection against influenza in humans (9). Antibody maturation selects for specificity over cross-reactivity of B cell-mediated immunity (i.e., affinity maturation), which may limit the breadth of female-biased immunity and protection when faced with rapid mutations associated with an RNA virus, such as influenza. Using viruses with mutations in antigenic sites of the HA protein of mouse-adapted A/California/04/2009 H1N1 (maA/Cal/09) (20), we tested the hypothesis that female-biased immunity and protection would be dependent on the extent of virus diversity as well as class switch recombination and somatic hypermutation mechanisms in B cells which constrains the breadth of epitope recognition. Vaccinated female mice benefit from the greater production of class-switched, somatically-hypermutated antibody by germinal center (GC) B cells to recognize diverse maA/Cal/09 antigens and be protected against influenza infection and disease; there are, however, limits to the female-biased immunity and protection that are dependent on the extent of virus mutations.

## Materials and Methods

### Mice

Male and female C57BL/6CR mice (7-8 weeks of age) were purchased from Charles River Laboratories (Frederick, MD). Male and female wild-type (WT), *mu*MT, and *Rag1* knock-out (*Rag1-/-*) mice on a C57BL/6J background (7-8 weeks of age) were purchased from the Jackson Laboratories (Bar Harbor, ME). The *Aicda* (activation-induced cytidine deaminase) knockout mice on a C57BL/6J background were maintained at the National Institute on Aging. Mice were housed at a maximum of 5 mice per cage under standard biosafety level 2 housing conditions in the Johns Hopkins Bloomberg School of Public Health. Food and water were available ad libitum. All animal procedures were approved by the Johns Hopkins University Animal Care and Use Committee under animal protocol MO18H250.

### Recombinant Virus Generation

The maA/Cal/09 challenge viruses contain mutations in the major antigenic regions of the HA head in the following positions: one mutation (1M) at K180Q; two mutations (2M) at K180Q and G157E; three mutations (3M) at K180Q, G157E, and N211D; and a substitution (Sub) at K180Q with the entire Ca2 antigenic region substituted with a non-human H5 sequence (**Supplementary Table 1**). Recombinant viruses were generated using the IAV 12-plasmids reverse genetics system (22, 23). Recombinant maA/Cal/09 and 1M viruses were generated previously (20). HA escape mutations G157E on the Ca2 antigenic site, the N211D on Sb antigenic site, and the Ca2 sequence substitution with H5 A/Vietnam/1203/2004 (24, 25) were introduced in pHH21-HA-maA/California/4/2009-K180Q to generate pHH21-HA to make recombinant 2M, 3M, or Sub viruses by either site directed mutagenesis (QuikChange Lightning Site-Directed Mutagenesis Kit, Agilent Technologies) or DNA synthesis (GenScript) (**Supplementary Table 1**). Eight maA/Cal/09 viral genome pHH21 plasmids with four helper plasmids expressing A/Udorn/72 (H3N2) PA, PB1, PB2, and NP proteins (23) were transfected into 50% confluent HEK293T cells in 6-well plates with TransIT-LT1 transfection reagent (Mirus Bio) according to the manufacturer’s protocol. After 24 hours of incubation at 37°C with 5% CO_2_, 4 µL of 5 mg/mL of *N*-acetyl trypsin were added in each 6-well for 4 hours at 37°C/5% CO_2_. Madin-Darby Canine Kidney cells (MDCKs, ∼5 × 10^5^ cells/ well) with 20 µL/well of 30% bovine serum albumin (BSA) were then added and kept at 37°C/5% CO_2_. Supernatants from the culture (1 mL) were collected and replaced with fresh infectious media with 5 µg/mL of *N*-acetyl trypsin daily for 7 days and then titrated using 50% tissue culture infectious dose (TCID_50_). Virus positive collection at the earliest time point was purified using plaque assays on MDCK cells in 6-well plates. Plaque-purified viruses were expanded to seed stocks. The HA sequences of the seed stocks were confirmed using RT-PCR of viral RNA and followed by Sanger sequencing. Seed stocks were then expanded into working stocks for experiments.

### Plaque Assays

To grow virus plaques, MDCK cells were grown to 100% confluency in complete medium (CM; DMEM, fetal bovine serum (FBS), L-glutamine, and Pen-strep) in 6-well plates as we have published before (23, 26, 27). The CM was removed, cells were washed with phosphate buffered saline (PBS)+ (1xPBS with 2mm calcium and magnesium) twice, and 400µL of virus inoculum was added to cells. Plates of cells were incubated at 32°C for 1 hour with rocking. The virus inoculum was removed after 1 hour of incubation and phenol-red free DMEM supplemented with 3% BSA, 100U/ml pen/strep, 2mM Glutamax (Gibco), 5µg/ml *N*-acetyl trypsin, and 1% agarose was added. Plates of cells were then incubated for 3-5 days at 32°C and fixed with 4% formaldehyde. The topmost agarose was removed, and cells were stained with napthol-blue black (Sigma Aldrich). Plaque size was analyzed in Image J (28). For the 1M, 2M, 3M, and Sub recombinant virus production, virus plaques were picked with a pipette and placed in media and stored at -80°C for ultimate generation of virus seed stocks. To calculate plaque forming units (PFU/mL) for each virus, the number of plaques were counted and divided by the given dilution used to seed virus.

### Low-MOI Virus Growth Curves

Low-MOI (multiplicity of infection) growth curves were performed as we have previously published, (23, 27),at a MOI of 0.001 on MDCK cells, which were infected in infectious media (IM; DMEM, 10% BSA, 5% L-glutamine, 5% pen-strep, and 5µg/mL *N*-acetyl trypsin) at 32°C for 1 hour. After infection, the virus inoculum was collected, cells were washed three times with 1xPBS+, new IM was added, and cells were placed at 32°C. At 1-, 12-, 24-, 36-, 48-, 72-, and 96-hours post-inoculation, IM was removed from the MDCK cells and stored for TCID_50_ infectious virus quantification. After each time point, fresh IM was added.

### Generation of Vaccine and ELISA Proteins

The maA/Cal/09 H1N1 virus (generated from available sequences by Dr. Andrew Pekosz (29)) was maintained and grown in IM on MDCK cells. To generate vaccine, maA/Cal/09 virus was grown at an MOI of 0.01 in MDCK cells for three days, upon which virus supernatant was collected, centrifuged, and inactivated with the addition of 0.05% β-propiolactone for 24 hours followed by a 2-hour 37°C incubation. Heat and chemically inactivated virus was purified by ultracentrifugation in a in a Beckman SW28TU rotor at 25,000rpm for 1 hour at 4°C with a 20% sucrose gradient and the virus pellet was resuspended in 1xPBS+. To confirm virus had been successfully inactivated, a TCID_50_ assay was performed. ELISA protein for maA/Cal/09, 1M, 2M, 3M, and Sub was grown similarly to vaccine except no heat or chemical inactivation steps were performed. A bicinchoninic acid assay (BCA, Pierce) was used to estimate the relative viral protein concentration.

### Mouse Lethal Dose 50 Assays (mLD_50_)

Groups of immunologically naïve mice were anesthetized with a ketamine/xylazine cocktail (80mg/kg and 5mg/kg) and inoculated intranasally with either 10^1^, 10^2^, 10^3^, 10^4^, 10^5^, or 10^6^ TCID_50_ of the maA/Cal/09, 1M, 2M, 3M, and Sub viruses. Survival was monitored for 21 days and the mLD_50_ for each sex for each virus was calculated via the Reed-Muench method (21, 30). If mice lost 30% or more of their mass, then they were humanely euthanized.

### Mouse Vaccination, Challenge, and Morbidity

Mice received an intramuscular injection of 20μg of maA/Cal/09 inactivated vaccine in 40µL of 1xPBS in the right thigh muscle for the prime (day 0) and boost (day 21) vaccinations. For live virus challenge, mice were anesthetized with a ketamine/xylazine cocktail (see above) and inoculated intranasally 6 weeks after vaccine prime (42 days post vaccination) with 10^5^ TCID_50_ of the 1M, 2M, or Sub virus suspended in 30μL of DMEM. As a correlate for morbidity, changes in mouse body mass were measured daily following live viral challenge.

### Virus Quantification in Mouse Lungs

Lungs were collected from mice at several timepoints post live viral challenge. Lungs were homogenized and serially diluted in IM before being plated on a monolayer of 100% confluent MDCK cells in replicates of 6. Cells were left to incubate for 6 days at 32°C and 5% CO_2_. Cells on plates were fixed for 1 hour in 4% formaldehyde and then stained with naphthol blue black for at least 4 hours. Plates were scored for cytopathic effect and the Reed-Muench method (30) was used to calculate the TCID_50_ value in the lungs for each animal.

### Enzyme-Linked Immunosorbent Assays (ELISAs)

Anti-maA/Cal/09, 1M, 2M, 3M, or Sub H1N1 IgG and IgG2c antibody titers were measured in plasma as described previously (20). ELISA plates (Microlon 96 well high binding plates; Greiner Bio-One) were coated with 100ng of purified virus (see above) for 24 hours at 4°C in carbonate bicarbonate buffer (pH 9.6). Plates were washed 3 times with 1xPBST (1xPBS +0.1% Tween-20) and blocked for 1 hour at 37°C with 10% nonfat milk powder in 1xPBS. After 1 hour, plasma samples were serially diluted in wash buffer plus 10% BSA and nonfat milk powder and added to the plates for 1 hour at 37°C. After washing 3 times, secondary antibodies for either anti-mouse horseradish peroxidase (HRP)-conjugated IgG (1:250, Invitrogen), IgM (1:3,000, Invitrogen), or IgG2c (1:20,000, Thermo Fisher Scientific) were added for 1 hour at 37°C. After 3 washes with 1xPBST, reactions were developed with 3,3’,5,5’ tetramethylbenzidine (TMB, BD Biosciences) and stopped with 1N hydrochloric acid (HCl) after 20 minutes. The absorbance was read at 450nm, and titers were calculated as the highest serum dilution with an average OD value above 3 times the average OD of the negative controls.

### Microneutralization Assays

The anti-maA/Cal/09, 1M, 2M, 3M, and Sub H1N1 neutralizing antibody (nAb) response in plasma was determined in tissue culture micro-neutralizing assays with MDCK cells as described previously (20). Plasma was heat inactivated for 35 minutes at 57°C before use. Plasma was diluted two-fold in IM and combined with 100 TCID_50_ of the maA/Cal/09, 1M, 2M, 3M, or Sub H1N1 virus for 1 hour at room temperature. The combined plasma and virus were added in duplicate to confluent MDCK cells for 24 hours at 32°C. The next day, plates were washed once with 1xPBS, and new IM was added. Plates were incubated at 32°C for 6 Days. Plates were fixed with 4% formaldehyde for 1 hour and stained with naphthol blue black for 4 hours. The nAb titer was calculated as the highest serum dilution that prevented cell death in half of the wells.

### Phage ImmunoPrecipitation Sequencing (PhIP-Seq) and Analysis

PhIP-Seq, using mouse serum, was performed as described in the published protocol (31). Briefly, 20μL of 1:100 diluted plasma in 1xPBS from individual mice were mixed with a 56 amino acid peptide library that tiles across the proteins of all human viruses (VirScan) (32), then immunoprecipitated using protein A and protein G coated magnetic beads. A set of 8 “mock” immunoprecipitations (IPs, i.e., no serum input) was run on the same plate. Beads were subsequently washed and resuspended in PCR master mix. 20 cycles of PCR were performed, followed by 20 more cycles of PCR with sequencing adapter and sample barcode containing primers. Amplicons were pooled and then sequenced on an Illumina instrument using a 1×50 cycle protocol.

Sequencing reads were mapped to the VirScan library using perfect matching. By comparing against the set of mock IPs, a fold change and differential abundance statistic were calculated for each peptide in each sample using the edgeR bioconductor package (33). Peptides were considered “hits” for a sample if the following conditions were met: count ≥ 15, p-value ≤ 0.001, and fold-change ≥ 5. Hits fold-change (HFC) values report the fold-changes of hits and is set to 1 for non-hits. Only the HA peptides from IAVs were considered in this study.

### Flow Cytometry and FACS

The numbers of GC B cells in the spleen were estimated using flow cytometry as previously described (20). Spleens were removed at 35 days post vaccination and a single cell suspension was made in FACS buffer (1xPBS, 1% heat inactivated FBS, 25mM HEPES, and 1mM EDTA). Spleen suspensions were filtered twice through a 70*µ*m filter. Red blood cell lysis was performed using ACK lysis buffer for one minute. The Fc receptors in each sample were blocked using anti-CD16/32 (BD Biosciences) for 20 minutes at 4°C. The total number of live cells were counted using a Cellometer Auto 2000 (Nexcelom Bioscience) with an AOPI ViaStain solution (Nexcelom Bioscience). After incubation, an antibody cocktail for GC B cells (B220+CD38-GL7+) was added and incubated for 20 minutes at 4°C in the dark. Antibodies included rat anti-mouse PE-Cy7-conjugated CD45R/B220+ (BD Biosciences), rat anti-mouse FITC-conjugated GL7 (BD Biosciences), and rat anti-mouse BV421 CD38 (BD Biosciences). Cell numbers were acquired using either FACSAria (BD Biosciences) or a Mo Flo XPD High Speed Cell Sorter (Beckman Coulter) and analyzed using FlowJo v.10 (Tree Star, Inc.). GC B cells were sorted using a BD FACS Aria Fusion instrument (BD Biosciences).

### Immunofluorescence and Confocal Microscopy

Spleens collected from male and female mice at 35 days post vaccination were fixed with 4% paraformaldehyde. Samples were mixed with 10% and then 30% sucrose to remove water and embedded in OTC and frozen at -80°C. Serial 10µm thick sections were taken from the frozen tissue block using the cryotome and slices were fixed onto slides with cold acetone for 10 minutes. Slides were washed twice with 1xPBS and incubated in 0.3% H_2_O_2_ solution for 10 minutes. After blocking with 0.1% titronX-100 in 5% BSA for 1 hour at room temperature, a PNA-biotinylated antibody (Vector Bio) was applied at 4°C overnight. Slides were washed with 1xPBS and incubated with Alexa Fluor 488 streptavidin (Biolegend) as a secondary antibody and Alexa Fluor 647 anti-mouse IgD (Biolegend) for 1 hour at room temperature. Sections were washed one final time and mounted with a DAPI solution and covered with glass and stored at -80°C.

Confocal fluorescent images were acquired on a Zeiss Axio Observer.Z1 (Zeiss) fluorescent microscope with Colibri.2 LED light source and an ORCA-R2 digital CCD camera (Hamamatsu) using ZEN imaging software. Tiling and stitching function was used to generate high-resolution images that covered entire spleen sections in a continuous field using a 5X objective. For each of the 12 male and 13 female spleens dissected and mounted, 3 spleen sections were imaged. Individual GCs were imaged at 10X magnification, and the relative area of each GC was calculated by measuring the quotient of the GC area (PNA+, stained green in images) over the entire B cell area (IgD+, stained magenta in images) using Fiji software (ImageJ). The relative GC area for 36 GCs from males and 38 GCs from female mice were measured.

### B cell isolation for real-time RT-PCR

Vaccinated and mock vaccinated male and female C57BL/6CR mice were euthanized at 35 days post vaccination and spleens were homogenized into a single cell suspension with red blood cell lysis. All CD19+ B cells were isolated (STEMCELL Technologies EasySep Mouse B Cell Isolation kit, STEMCELL Technologies) and mRNA was extracted and purified (PureLink RNA Mini Kit, Invitrogen). This mRNA was used to generate cDNA via RT-PCR. *Aicda* expression was measured by qPCR (Primers from Integrated DNA Technologies, Mm.PT.58.42247522). The relative gene expression was normalized to a *Gapdh* house-keeping gene and mock-vaccinated animals using the ΔΔCT method as described previously (20).

### Somatic Hypermutation Intron Sequencing and Analysis

Splenic B cells were isolated at 35 days post vaccination from male and female C57BL/6CR mice and GC B cells (B220+CD38-GL7+) were sorted as described above. Cells were then lysed in digestion buffer (10 mM Tris, pH 8.0, 25 mM EDTA, 100 mM NaCl, 1% SDS, and 0.1 mg/ml proteinase K) at 55°C overnight. Genomic DNA was isolated by phenol/chloroform extraction and ethanol precipitation. The J_H_4 intronic region was amplified using a nested PCR protocol with Herculase II high-fidelity polymerase (Agilent). Primary PCR was performed for 25-cycles using V-region forward primer (5’-AGCCTGACATCTGAGGAC-3’) and intron reverse primer (5’-GAGCCTCACTCCCATTCCTCGG-3’), followed by a 35-cycle secondary PCR using V-region primer (5’-GCCTGACATCTGAGGACTCTGC-3’) and intron reverse primer (5’-TAGATGCCTTTCTCCCTTGACTCA-3’). The 492 bp of J_H_4 intronic DNA was sequenced and unique VDJ clones were analyzed for mutations.

### Statistical Analyses

Graphs were created and statistics were performed in GraphPad Prism 9. T-tests (to compare between two groups), one-way ANOVA (to compare among three or more groups with single independent variable) and two-way ANOVA (to compare between groups when having two variables) were used to analyze plaque sizes, IgM/IgG/IgG2c titers, nAb titers, virus titers, changes in body mass, cell numbers, GC sizes, mutation frequencies, and gene expression data. For one and two-way ANOVAs, analysis was followed by Tukey’s multiple comparison tests. Differences were considered significant if *p*≤0.05.

## Results

### HA mutations in maA/Cal/09 viruses reduce virus growth *in vitro* and cause greater morbidity in naïve females than males *in vivo*

To test the breadth of female-biased antibody-mediated immunity against antigenically diverse viruses, naturally occurring mutations observed in human A/Cal/09 (24, 25) antigenic sites were introduced into the HA sequence of maA/Cal/09: 1) one point mutation (1M) in Sa; 2) two point mutations (2M) in Ca2 and Sa; 3) three point mutations (3M) in Ca2, Sa, and Sb; and 4) a point mutation in Sa along with an antigenic site substitution of the Ca2 epitope with one derived from an H5 sequence (Sub) (**Supplemental Table 1, Figure 1A**). The overall plaque morphology of the parental maA/Cal/09 virus included medium-sized plaques with complete clearance (i.e., cell death) within the middle of the plaque. While the 1M and Sub viruses had similar plaque morphology as maA/Cal/09, the 2M and 3M virus plaques were significantly smaller in size (**Figure 1B-C**). Low MOI growth curves showed that the parental maA/Cal/09 virus grew significantly better by 48 hours post-infection (which corresponded to peak titer) as compared to the 2M and 3M viruses (**Figure 1D**). The mutant maA/Cal/09 viruses were each able to replicate and be released by MDCK cells, albeit to different peak titers and with different plaque morphologies.

**Figure 1:**
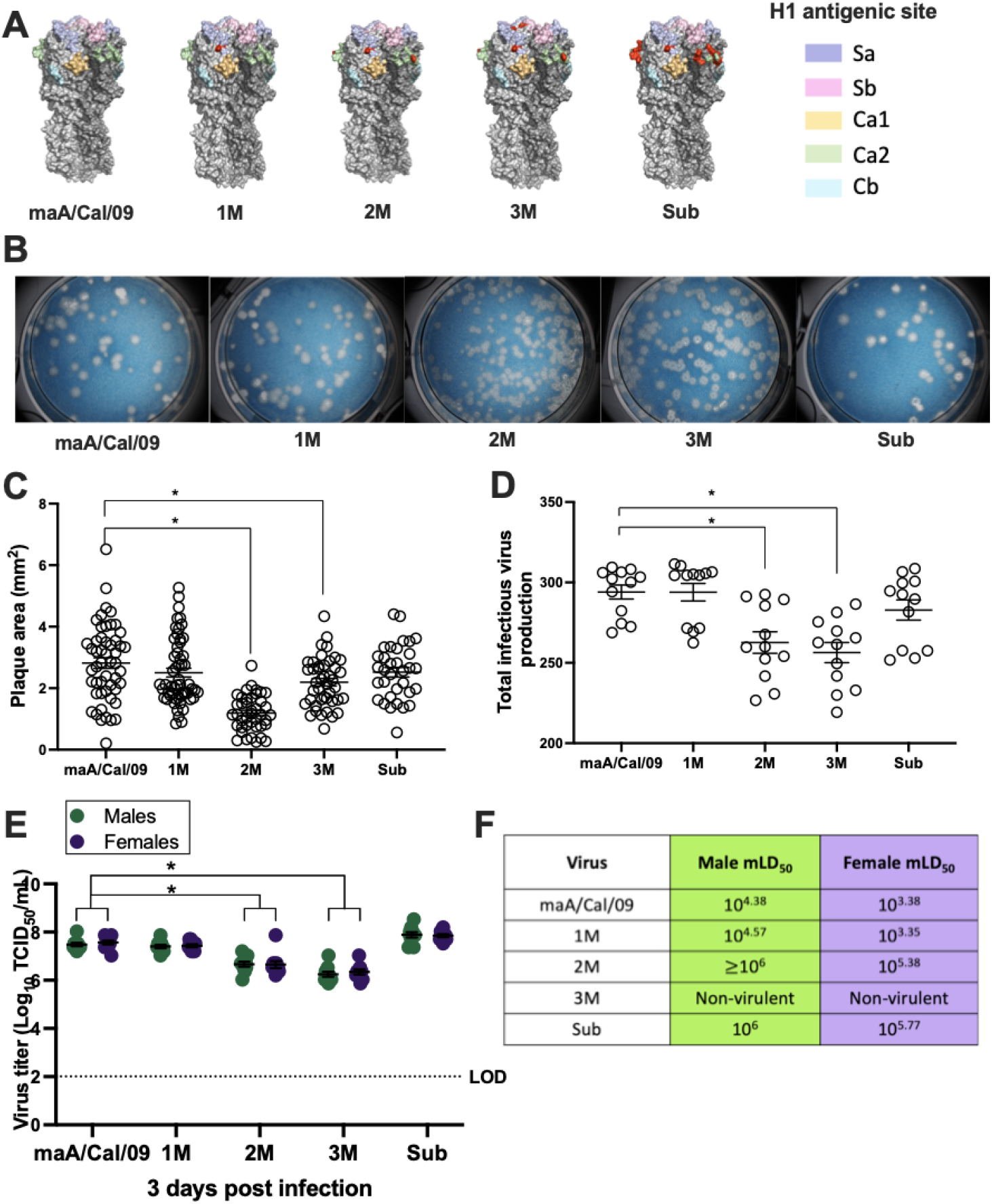
Mutations in mouse-adapted A/California/4/2009 H1N1 viruses cause differences in growth morphology and kinetics *in vitro* and sex differences in virulence *in vivo*. A) Mutations were inserted into antigenic sites of the parental mouse-adapted A/California/4/2009 (maA/Cal/09) H1N1 virus hemagglutinin (HA). Single point mutations were inserted into the Sa site (single mutant, 1M); Sa and Ca2 sites (double mutant, 2M); and Sa, Ca2, and Sb sites (triple mutant, 3M). A fourth virus had a single point mutation in the Sa site at the same position as 1M but had a non-human, avian H5 sequence substituted into the Ca2 region (Sub). The H1 major antigenic sites are shown in pastel shades on the head of the HA protein trimers (different shades of gray). Mutations are shown in red. The HA structures were created in *PyMOL* with PDB: 3LZG for A/California/4/2009. B) Viruses were used to perform plaque assays to determine growth and plaque morphology on a monolayer of MDCK cells. C) Plaque sizes for each virus were measured and are plotted as area per mm^2^. D) Low multiplicity of infection (MOI) growth curves were used to compare the growth kinetics of the parental maA/Cal/09 virus and each of the mutant viruses, with area under the curve (AUC) calculated up to 48 hours post infection (i.e., peak titers). E) Lungs were collected from naïve (unvaccinated) male and female C57BL/6CR mice 3 days after infection with 10^5^ TCID_50_ units of each mutant virus and used to measure replicating virus (n=9-10/sex/virus). F) Log_10_ dilutions of each virus ranging from 10 – 10^6^ TCID_50_ units were used to intranasally infect male and female mice (n= 5-10/sex/virus dose) to calculate the mouse lethal dose 50 (mLD_50_). *Statistically significant difference at *p* < 0.05.

To assess the virulence of the mutant viruses, naïve male and female mice were intranasally infected with increasing doses (10^1^-10^6^ TCID_50_ units) of either the parental maA/Cal/09 virus or one of the four mutant viruses and either euthanized 3 days post infection to measure pulmonary virus titers (from mice infected with 10^5^ TCID_50_ units only) or followed for 21 days to quantify morbidity (i.e., body mass loss) and mortality. Following infection with 10^5^ TCID_50_ units of each virus, naïve males and females exhibited similar pulmonary titers of all infecting viruses (**Figure 1E**). The parental maA/Cal/09, 1M, and Sub viruses replicated to higher titers in the pulmonary tissue than the 2M or 3M viruses (**Figure 1E**). Dose-dependent survival curves were used to calculate the mLD_50_ value for each virus separately for males and females (**Figure 1F**). For the maA/Cal/09, 1M, 2M, and to a lesser extent the Sub virus, naïve female mice suffered greater mortality than male mice, with females having an mLD_50_ that was approximately one log lower than males (**Figure 1F)**. The 3M virus did not cause mortality at any dose in either sex of mice and was labeled as non-virulent. These data illustrate that as these viruses acquired more mutations, becoming antigenically more distinct from the parental virus, virulence and sex differences in mortality were reduced among naïve mice.

### Vaccinated females have greater antibody cross-reactivity against mutant maA/Cal/09 viruses than males

Previous studies have shown that female mice produce greater anti-mA/Cal/09 IgG and IgG2c, but not IgG1, titers than male mice following vaccination (16, 20). To determine if sex differences were observed in antibody recognition of mutant viruses, mice were vaccinated and boosted with maA/Cal/09 and then bled at 28 days post vaccination to measure cross-reactive total IgG and IgG2c antibodies against maA/Cal/09, 1M, 2M, 3M, and Sub viruses. Vaccinated females produced greater anti-maA/Cal/09 IgG (**Figure 2A**) and IgG2c (**Figure 2B**) titers than males and had greater cross-reactive IgG (**Figure 2A**) and IgG2c (**Figure 2B**) against the 1M, 2M, 3M, and Sub viruses than males. Titers of cross-reactive IgG and IgG2c against the mutant viruses were similar to titers against the parental virus for both males and females (**Figure 2A-B**).

**Figure 2:**
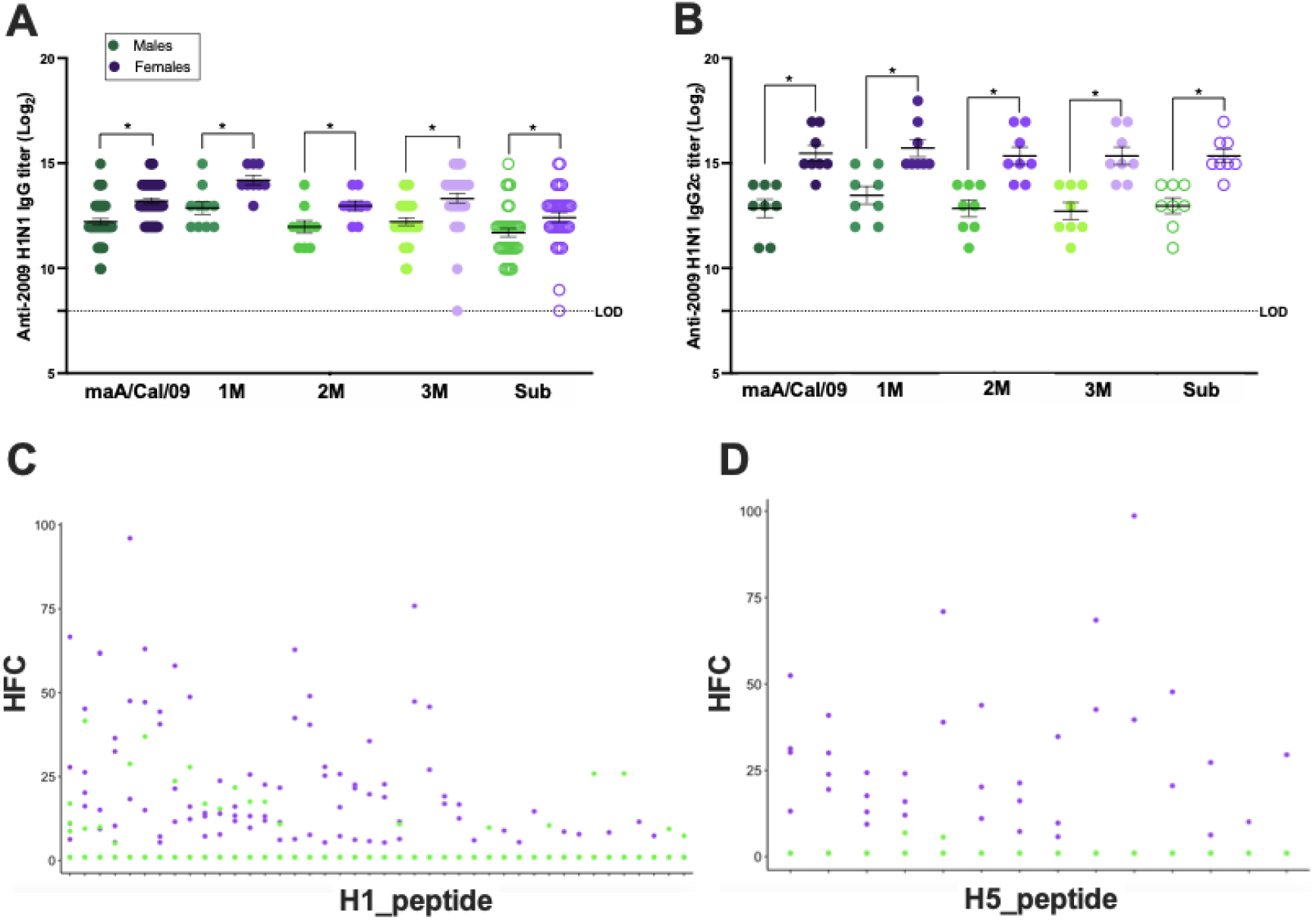
Vaccinated female mice mount greater cross-reactive antibody responses to mutant mouse-adapted A/California/4/2009 H1N1 viruses than males. Male and female C57BL/6CR mice were vaccinated and boosted with mouse-adapted A/California/4/2009 (maA/Cal/09) H1N1 virus and bled at 28 days post vaccination. Serum was used to measure cross-reactive IgG (n=9-45/sex) (A) and IgG2c (n=8/sex) (B) antibodies towards the vaccine virus maA/Cal/09 as well as the mutant viruses 1M, 2M, 3M, and Sub by ELISA. Serum from vaccinated males and females was used to determine antibody reactivity to thousands of linear influenza hemagglutinin epitopes via PhIP-Seq using the Vir-Scan library. Reactivity of these vaccine-induced antibodies to unique H1 epitopes (C) and cross-reactivity to H5 epitopes (D) are shown (n=9-10/sex), with each unique epitope represented on the X-axis and the hits fold-change (HFC) on the Y-axis. HFC is a measure of bound peptide from the serum-immunoprecipitated samples relative to the mock-immunoprecipitated samples, with no significant difference set to HFC = 1 (Methods). *Statistically significant difference at *p* < 0.05.

Greater breadth of recognition of HA viral epitopes might explain female-biased cross-reactivity against mutant maA/Cal/09 viruses. To evaluate recognition of linear viral epitopes, PhIP-Seq was performed using the VirScan library, whereby serum from vaccinated male and female mice collected either 28- or 35-days post vaccination was combined with a library of linear epitopes from IAV proteins. In total, 75 unique, linear HA epitopes were recognized by the vaccine-induced antibodies produced in at least 1/10 males or 1/10 females, with antibodies recognizing 42 H1 epitopes, 4 2009 H1 epitopes, 11 H2 epitopes, 14 H5 epitopes, and 8 H3 epitopes (**Supplemental Figure 1A**). Each of these 75 epitopes mapped to the HA2 or stem domain, where linear epitopes are most abundant. As an example, the binding footprint for the 4 2009 H1 epitopes (3 of which are unique) is illustrated in **Supplemental Figure 1B**. In each case, the percentage of female mice with reactivity to at least one HA epitope was greater than that for male mice (**Supplemental Figure 1A**). Not only was the percentage of vaccinated female mice that recognized unique H1 epitopes greater than for males, but the hits fold-change (HFC) values, which are a measure of how bound peptide from the serum-immunoprecipitated samples relative to the mock-immunoprecipitated samples, was greater for females compared to males for H1 epitopes (**Figure 2C**), H5 epitopes (**Figure 2D**), and H2 epitopes (**Supplemental Figure 1C**), which are all Group 1 IAVs. Serum from vaccinated females also was significantly better at recognizing H3 epitopes (**Supplemental Figure 1D**), which is a Group 2 IAV, providing evidence of greater potential for heterologous immunity in vaccinated females than males, which is consistent with previous studies utilizing live H1N1 and H3N2 viruses (21). These data suggest that because females have greater cross-reactivity of antibodies against novel HA epitopes, vaccinated females may have greater breadth of immunity against IAVs than males.

### Vaccinated females have greater cross-protection against infection and disease than males following challenge with mutant viruses, which is dependent on B cells

Neutralizing antibodies, which recognize primarily conformational rather than linear viral epitopes (12, 34), were measured as a correlate of protection. Although vaccinated females had greater nAb titers than males against the maA/Cal/09 vaccine virus, as demonstrated previously (20), nAb titers against the mutant maA/Cal/09 viruses were significantly reduced and sex differences were not apparent (**Figure 3A**). If live virus neutralization is the correlate of vaccine-induced protection, then females vaccinated against maA/Cal/09 may not be better protected against mutant maA/Cal/09 viruses. To test this hypothesis, vaccinated males and females were challenged with lethal doses of each of the virulent mutant viruses (1M, 2M, and Sub) at 42 days post vaccination. Subsets of mice were euthanized at 3 days post challenge to measure peak pulmonary virus titers while other mice were followed for 14 days for body mass loss as a measure of morbidity. Following live virus challenge, vaccinated female mice had lower pulmonary titers of the 1M and 2M, but not the Sub, virus than vaccinated males, with 60% (3/5) and 80% (4/5) of female mice clearing the 1M and 2M virus, respectively, compared with none of the male mice clearing virus at this timepoint **(Figure 3B**). Body mass loss over the 14 days post challenge was translated into area under the curve (AUC) to show individual data points, with greater mass loss indicated by lower AUC values. Male mice challenged with either the 1M or 2M virus experienced greater body mass loss (i.e., had lower AUC values) as compared with their female counterparts (**Figure 3C**). In contrast, challenge with the Sub virus did not cause sex differences in morbidity among vaccinated mice. Despite causing greater overall levels of virus replication in the lungs **(Figure 3B**), the Sub virus caused minimal disease in vaccinated male and female mice (**Figure 3C**).

**Figure 3:**
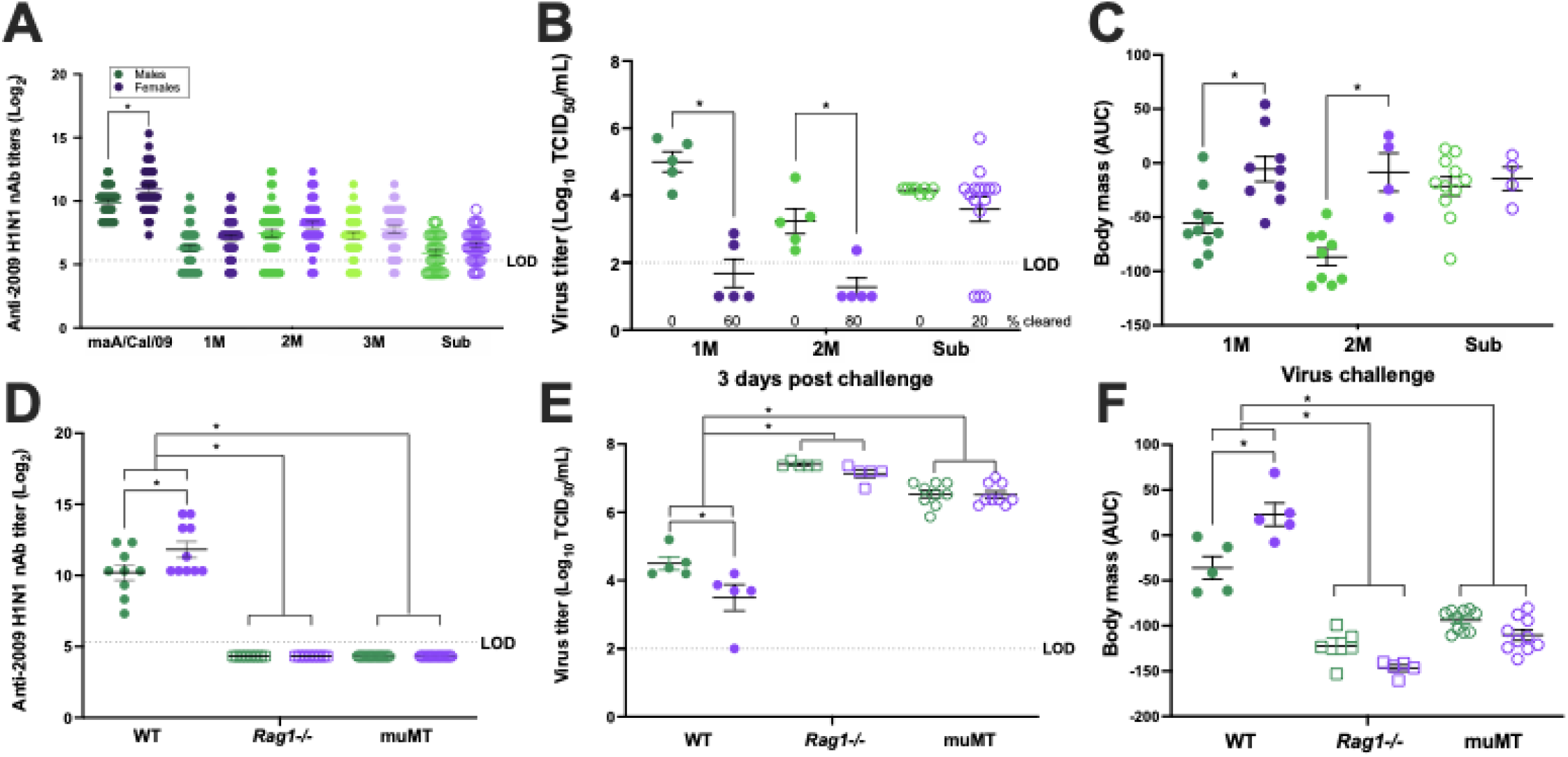
Female-biased, vaccine-induced protection from disease and virus replication following live virus challenge is dependent upon B cells. Male and female mice were vaccinated and boosted with mouse-adapted A/California/4/2009 (maA/Cal/09) H1N1 virus and challenged with 10^5^ TCID_50_ of the virulent mutant viruses 1M, 2M, or Sub. A) Neutralizing antibodies (nAb) towards each of the five viruses were measured in serum samples collected at 28 days post vaccination from wild-type (WT) C57BL/6CR mice (n=33-45/sex). B) Subset of WT male and female mice were euthanized at 3 days post-challenge to measure replicating virus titers in the lungs (n=5-15/sex). C) Body mass loss after virus challenge was tracked as a correlate for morbidity for 14 days and converted into area under the curve (AUC) values, with lower values indicating greater mass loss (n=4-11/sex). Male and female WT, *Rag1-/-*, and muMT mice on a C57BL/6J background were vaccinated and boosted with mouse-adapted A/California/4/2009 (maA/Cal/09) H1N1 virus. D) Serum from 28 days post vaccination was used to measure nAb titers to the maA/Cal/09 H1N1 vaccine virus in knock out compared with WT male and female mice (n=10-20/sex). Vaccinated and boosted WT, *Rag1-/-*, and muMT male and female mice were challenged with 10^5^ TCID_50_ of the 2M mutant virus and subsets of mice were used to measure replicating virus titers in the lungs at 3 days post-challenge (n=5-9/sex). F) After virus challenge, WT, *Rag1-/-*, and muMT male and female mice were monitored for body mass loss for 14 days, with lower AUC indicating greater mass loss (n=5-10/sex). *Statistically significant difference at *p* < 0.05.

Because live virus neutralization did not adequately predict the female-biased protection against mutant maA/Cal/09 viruses, we sought to ensure that vaccine-induced protection was indeed B cell and not T cell mediated. Male and female WT, *mu*MT (mature B cell deficient), and *Rag1-/-* (mature B and T cell deficient) mice were vaccinated and boosted with inactivated maA/Cal/09 and antibody responses as well as outcomes following challenge with live 2M virus were analyzed. After vaccination, nAb titers against the maA/Cal/09 vaccine virus were significantly greater among WT female than male mice (**Figure 3D**). In the absence of B cells, either alone or in combination with T cells, nAb titers were not detectable in either sex (**Figure 3D**). Live 2M virus challenge of vaccinated WT mice resulted in lower pulmonary virus titers and reduced body mass loss among females as compared with males (**Figure 3E-F**). In contrast, among both *mu*MT and *Rag1-/-* mice, 2M virus titers in the lungs as well as morbidity were greater than among WT mice and sex differences were no longer observed (**Figure 3E-F**). Vaccinated *mu*MT and *Rag1-/-* mice had similar outcomes following challenge with live 2M virus, suggesting that in the absence of B cells, with or without T cells, vaccine-induced, female-biased protection against challenge with a mutant maA/Cal/09 virus was eliminated.

### Splenic GC size and frequencies of B cells is greater in vaccinated female than male mice

Following vaccination with IIV, antibodies undergo affinity maturation to increase the overall specificity, functionality, and quantity of antibodies produced in GCs in lymph nodes (LNs) and the spleen (35, 36). Because vaccinated female mice produced more IgG and IgG2c following vaccination **(Figure 2A-B**) and experience greater protection from virus-induced infection and disease (**Figures 3B-C**), we hypothesized that females may have greater GC activity than males. Male and female mice were vaccinated and boosted with maA/Cal/09 and whole spleens were dissected at 35 days post boost. Spleens from vaccinated male and female mice were frozen, sectioned, and stained to outline total B cell areas (IgD+, PNA+) and GCs (IgD-, PNA+, **Figure 4A**). While the overall number of GCs in spleen sections from vaccinated male and female mice was not different (**Figure 4B**), the percentage of the GC area relative to the follicle area was significantly greater in spleens from females than males, indicating that the relative size of the GCs was larger in female than male mice (**Figure 4C**). Flow cytometry was used to quantify numbers of GC B cells in the spleens of vaccinated male and female mice. Both the percent of B220+ GC B cells (a marker for mature B cells in mice, **Figure 4D**), as well as the total number of GC B cells (**Figure 4F**) in the spleen were greater in vaccinated female than male mice. These data suggest that greater activation of GC B cells underlies greater antibody class switch recombination and possibly somatic hypermutation to result in greater cross-protection against mutated maA/Cal/09 viruses in vaccinated females than males.

**Figure 4:**
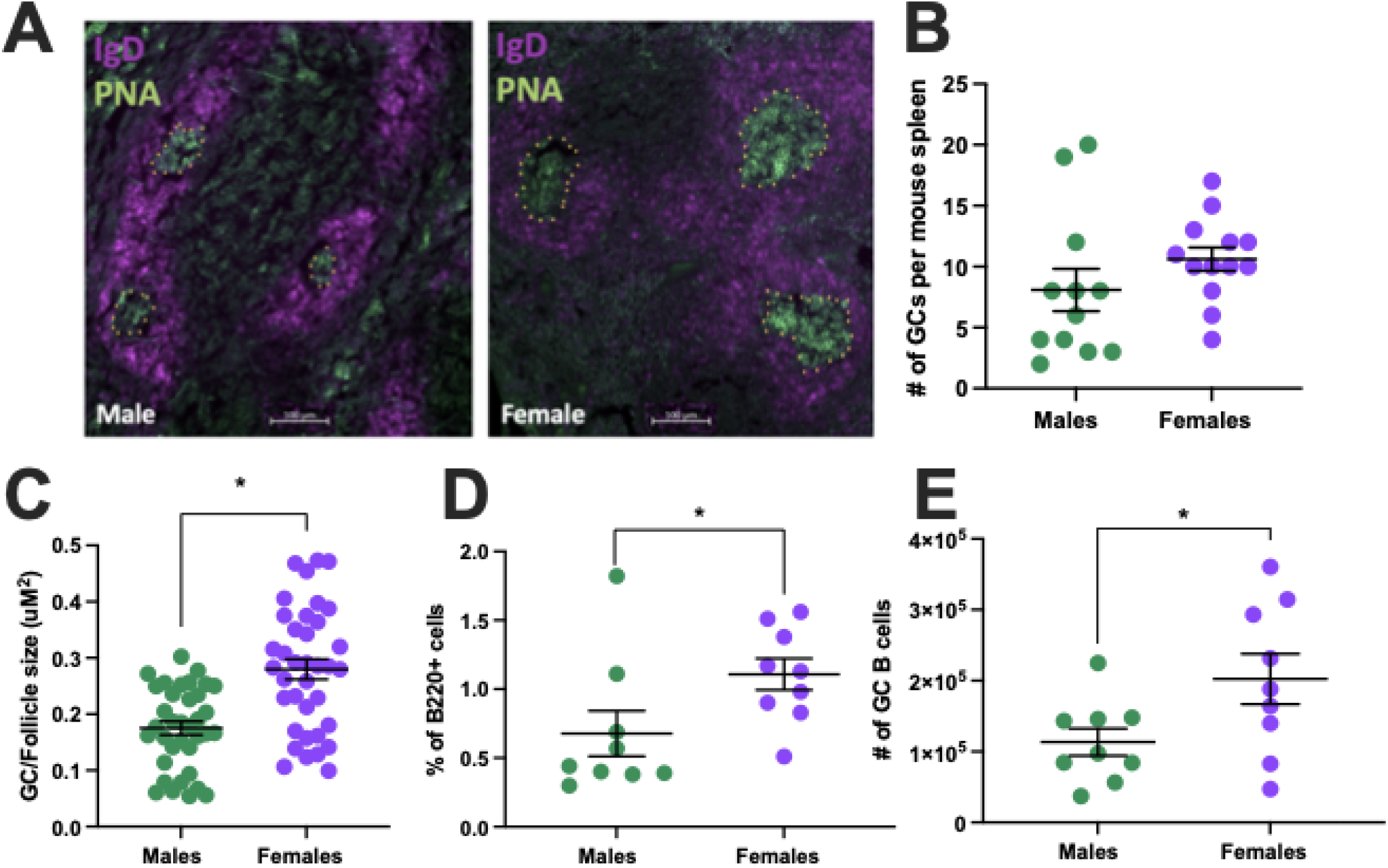
Influenza vaccination elicits a more robust germinal center B cell response in female than male mice. Male and female C57BL/6CR mice were vaccinated and boosted with mouse-adapted A/California/4/2009 (maA/Cal/09) H1N1. A) At 35 days post vaccination, mice were euthanized, and spleens were dissected for gross histology and microscopy at 10X magnification, where IgD^+^ cells stain magenta and PNA^+^ cells stain green. Germinal centers (GCs) are outlined in a yellow stippled line. B) The total number of GCs per spleen was quantified (n=12-13/sex) from 3 sections per animal. C) Relative size of GCs (IgD-, PNA+) relative to its surrounding follicle (IgD+, PNA-) was measured in 36 GCs from males and 38 GCs from females (n=5/sex). D) The percentage of B cells that were B220^+^ was measured via flow cytometry together with the total number of GC B cells in male and female mice after vaccination (n=9/sex) (E). *Statistically significant difference at *p* < 0.05.

### Increased antibody maturation underlies greater vaccine-induced protection against mutant maA/Cal/09 viruses in females than males

Because class-switched IgG antibody titers and frequencies of GC B cells were greater among vaccinated females than males, we next sought to determine the mechanism of antibody diversity that was responsible for the elevated antibody function in females compared with males after vaccination. At 35 days post vaccination, expression of *Aicda* mRNA (i.e., the gene that encodes for AID, which is a key enzyme in class switch recombination and somatic hypermutation antibody diversity reactions) was measured in B220+ B cells and was greater in B cells isolated from spleens of vaccinated female than male mice (**Figure 5A**). To test the hypothesis that sex differences in vaccine-induced immunity and protection against mutated maA/Cal/09 viruses were dependent on this elevated *Aicda* expression, WT and *Aicda-/-* male and female mice received the prime and boost maA/Cal/09 vaccine and were bled at 28 days post vaccination to measure antibody titers. Without the ability to undergo class switch recombination, sex differences in antibody responses were eliminated in *Aicda-/-* mice, with both sexes producing greater titers of IgM (**Figure 5B**) and lower titers of IgG (**Figure 5C**) than WT animals. Neutralizing capability towards the vaccine virus was significantly reduced in *Aicda-/-* compared to WT females, but not males (**Figure 5D**). Finally, in the absence of antibody diversity, vaccinated *Aicda-/-* mice were more susceptible to challenge with live 2M virus than WT mice. In the absence of AID, sex differences in vaccine-induced protection against pulmonary virus replication and morbidity (i.e., body mass loss) were eliminated, and the magnitude of this effect was greater for females than males when compared with WT mice (**Figure 5E-F**).

**Figure 5:**
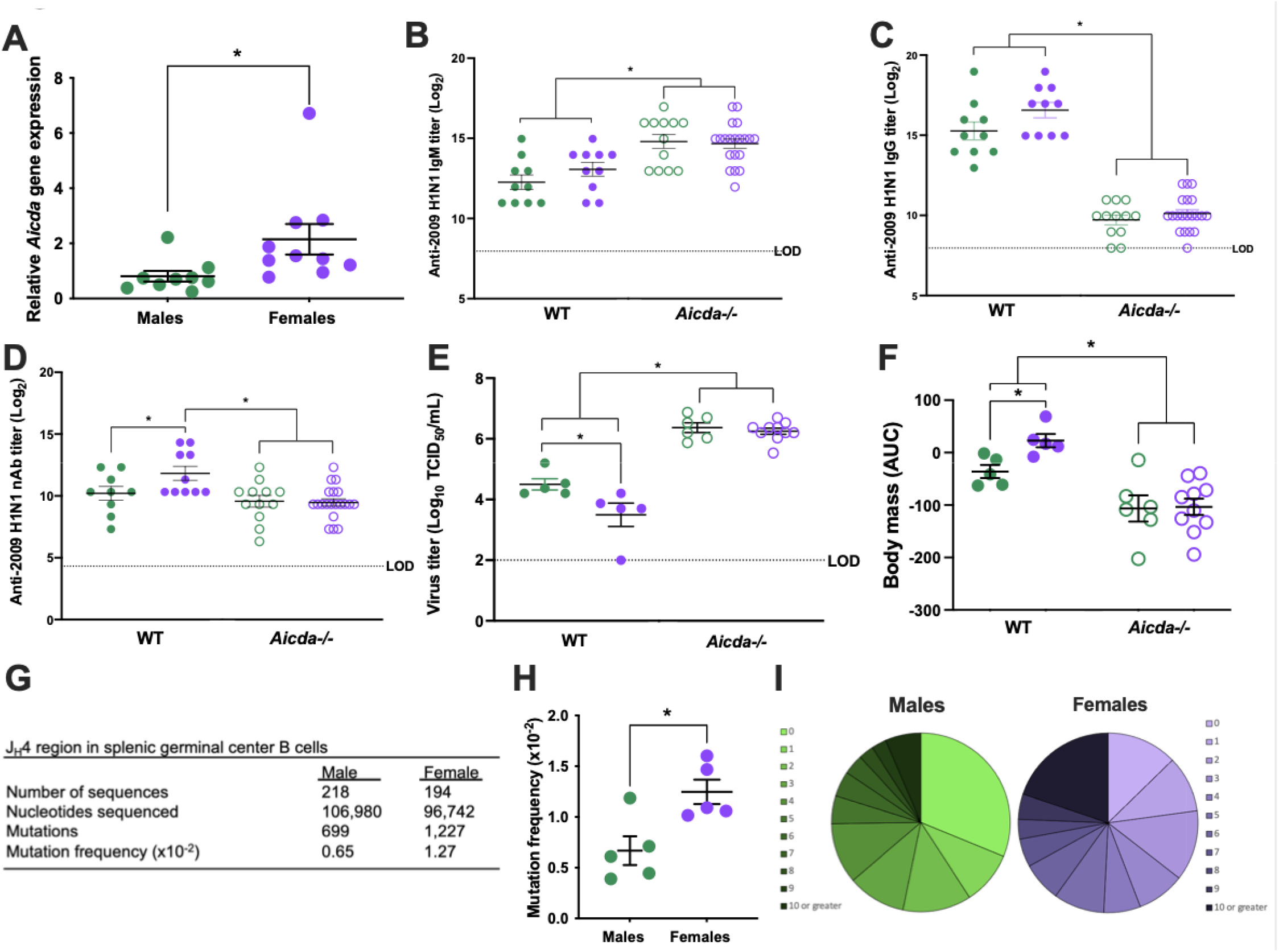
Greater dependence on and frequencies of somatic hypermutations in germinal center B cells mediates greater influenza vaccine-induced immunity and protection in female than male mice. Male and female C57BL/6CR were vaccinated and boosted with mouse-adapted A/California/4/2009 (maA/Cal/09) H1N1 vaccine virus. A) Splenic B cells from male and female mice were isolated at 35 days post vaccination to measure activation-induced cytidine deaminase (*Aicda)* gene expression (n=9-10/sex). Serum from vaccinated and boosted wild type (WT: C57BL/6J) and *AID-/-* male and female mice was (n=10-20/sex) used to measure anti-maA/Cal/09 IgM (B), IgG (C), and neutralizing antibody (D) titers. E) WT and *Aicda-/-* animals were challenged with 10^5^ TCID_50_ of the 2M mutant virus and males and females were euthanized (n=5-10/sex) at 3 days post challenge to measure replicating virus titers in the lungs. F) Vaccinated male and female mice (n=5-10) were challenged with 10^5^ TCID_50_ of the 2M virus and were monitored for body mass loss over 14 days, with data converted into area under the curve (AUC) values in which a lower AUC corresponds with greater mass loss. G) Pooled antibody somatic hypermutation in germinal center B cells after vaccination in the J_H_4-intronic region was analyzed (n=5/sex). H) The mutation frequency in the J_H_4-intronic region of antibodies from splenic germinal center B cells collected 35 days after vaccination and the percent of total sequences containing increasing numbers of mutations per sequence are shown as a gradation of darker color in the pie chart (n=5/sex) (I). *Statistically significant difference at *p* < 0.05.

During somatic hypermutation, point mutations are accumulated in the V gene sequences of the B cell receptor, a process that is regulated by AID and underlies antibody diversity (36). To determine if females had greater vaccine-induced somatic hypermutation, the J_H_4 intronic region of recombined V genes from GC B cells collected from spleens 35 days post vaccination were sequenced to compare mutation frequencies. The overall number of mutations and corresponding mutational frequency were significantly greater in sequences from vaccinated female than male mice (**Figure 5G-H**). Within the 218 male and 194 female regions sequenced, female mice had more mutations per sequence, with the majority displaying 10 or more mutations, whereas a greater percentage of male sequences were unmutated (**Figure 5I)**. These data suggest that highly mutated, class-switched antibodies afford females an advantage in terms of vaccine-induced protection against infection and disease after challenge with mutant maA/Cal/09 viruses.

## Discussion

Females develop greater antibody responses to influenza vaccines than males (9, 16, 17, 19-21, 37, 38), however, there are limits to female-biased protection following vaccination. Occasionally, the selected influenza vaccine strains have significant antigenic differences compared to circulating influenza viruses, which may be due to egg adaptation or other variables (5). In these vaccine mismatch years, such as the 2017-18 season in North America, the female-biased protection is lost (9). We sought to characterize these limits to female-biased antibody responses and protection after vaccination and found that antibodies from vaccinated female mice are better able to recognize diverse HA epitopes and protect against mutant H1N1 viruses than antibodies from males. Unlike the well accepted dogma that vaccine-induced nAb and IgG responses predict the degree of protection against influenza, our data show that this is not equally true for biological males and females. In females, heightened cross-recognition of mutant H1N1 viruses did not translate into heightened cross-neutralization to these viruses, as vaccinated male and female mice experienced equal neutralizing capabilities. In males, having equal nAb responses to mutant maA/Cal/09 viruses as compared to females did not predict equal protection. Rather, the greater cross-reactivity of vaccine-induced IgG antibodies to diverse HA epitopes successfully correlated with greater protection from severe disease and pulmonary virus replication in females after live virus challenge with H1N1 mutants.

Cross-protection against mutant H1N1 viruses among females compared with males was dependent on B cells as deletion of B cells either alone (*mu*MT mice) or in combination with T cells (*Rag1-/-* mice) eliminated female-biased protection following vaccination. Vaccination induced greater numbers and activity of GC B cells in females than males. This suggests that the presence of functional B cells is necessary for female-biased protection. We also studied post-vaccination GCs to understand how the GC reaction impacted female-biased protection. Our data show that vaccination induced greater numbers of GC B cells and greater expression of the somatic hypermutation- and class switch recombination-associated gene *Aicda* in female than male mice. Within these GC B cells, greater somatic hypermutation frequencies were seen following vaccination of female compared with male mice. In the absence of antibody diversity (*Aicda-/-* mice), the female-biased immunity and protection was eliminated, illustrating that functional GC B cells and greater antibody diversity is necessary for the female bias in cross-protection against mutant H1N1 viruses. We hypothesize that increased expression of *Aicda* and frequency of somatic hypermutations in GC B cells may in part be due to the numerous estrogen and progesterone response elements that have been mapped to the promoter region of the *Aicda* gene (39-42). Studies of antibody production in the context of autoimmune diseases reveal that estrogen regulation of *Aicda* is critical for female-biased antibody production and disease progression (41). While supplementation of B cell cultures with exogenous estrogen increases IgG heavy chain transcription due to the presence of estrogen receptors in immunoglobulin regulatory elements and switch sites (43, 44), whether estrogens or even progesterone can therefore directly regulate somatic hypermutations to improve vaccine-induced immunity is not known and is currently under investigation in our laboratory.

A limited number of studies have evaluated the impact of sex steroids on immune responses to vaccines. Estradiol, at physiological concentrations, can stimulate antibody production by B cells, including antibody responses to an IIV administered in mice (16, 45). In humans, reduced nAb responses to influenza vaccination are correlated with higher serum testosterone concentrations (17) and in mice, greater testosterone concentrations cause reduced antibody and CD8+ T cell activity following malaria vaccination (46). Elevated testosterone concentrations in males also are associated with greater lipid metabolism, suggesting that the immunosuppressive role of testosterone and the reduced antibody responses to vaccines in males may be mediated by the expression of genes involved in lipid metabolism that are associated with the suppression of inflammatory responses (17).

Some of the drawbacks to this study include the inability to produce mutant viruses for each of the five major H1 antigenic sites (Sa, Sb, Ca1, Ca2, and Cb). At the time of this study design, we were unable to find documented point mutations in the Ca1 and Cb regions that would abrogate antibody recognition and neutralization. Having more mutant viruses to diverse combinations of antigenic sites would also have allowed for us to estimate if there was a difference in antibody/epitope usage or hierarchy, a phenomenon which is known to be different after influenza vaccination versus infection (24, 47). Whether males and females induce antibodies or nAbs to influenza viruses in a differential hierarchical fashion is unknown. For this study, GC reactions from the spleen rather than the draining LNs were evaluated due to the greater cell numbers and size of the spleen. Further studies are required to compare the GC responses in spleens to LNs in mice after vaccination, as organ-specific GC responses to influenza infection have been observed (48). Whether antibody affinity and avidity to the mutant H1N1 viruses display a sex-differential phenotype was also not measured in the current study and tracking the progression of both in germline antibody as compared to post-vaccination antibody in male and female mice will be examined in future studies. Finally, interrogation into the role of CD4+ T cells, particularly T follicular helper cells (Tfhs), in conjunction with B cells and how they combined help with induction of GC B cell numbers and activity is necessary. This is especially important with regards to the formation of the differential GC response, as the cross talk between antigen, B cells, T cells, and antigen presenting cells (e.g., dendritic cells) is necessary for the formation of follicles and GCs. Further detailed exploration of this relationship, and the relationship with various hormonal stimuli, will be the basis for future studies. Interestingly, sex differences in some of the universal influenza vaccine platforms are not observed, which may be due to equal T cell help during GC formation resulting in similar overall antibody output in males and females (49-51).

One of the most well conserved immunological differences between the sexes is in antibody responses to foreign antigens (52). Beyond influenza vaccines, adult females have greater antibody responses to hepatitis B, yellow fever, rabies, herpes, and smallpox vaccines (52). Sex differences in antibody responses have evolved in diverse species. In birds, for example, females exhibit greater antibody responses to immune challenges and these effects are often most pronounced during the mating season when male testosterone concentrations are highest (53, 54). Why would sex differences in antibody responses evolve? We speculate that when offspring are young (i.e., during the neonatal period), protection against infection is primarily mediated by passive maternal immunity. In humans, a majority of maternal antibodies are transferred into fetal circulation, prior to birth, through the placenta (55). In some mammals, transfer of maternal antibodies can also occur through the colostrum immediately following birth and in milk for a longer duration after birth (55). Regardless of species, elevated production of antibodies caused by either vaccination or infection of females enhances the transfer of antibodies to the fetus or neonate to protect them during a critical period of infectious disease susceptibility. We postulate that natural selection favors increased antibody production, and the mechanisms therein, in females compared with males of reproductive ages because transfer of maternal antibodies from mother to young increases reproductive success by minimizing the detrimental effects of infection on offspring survival. While this might leave females more susceptible to antibody-mediated diseases, such as autoimmune diseases, these diseases typically cause morbidity after reproductive years and, therefore, would not be selected against. The next frontier will be harnessing these observations and data to improve vaccines and equitable protection in both males and females.

## Conflicts of interest

The authors have no conflicts to disclose.

## Author contributions

Rebecca Ursin, Robert Maul, Patricia Gearhart, Andrew Pekosz, and Sabra Klein conceived the experimental questions. Rebecca Ursin, Santosh Dhakal, Alice Mueller, and Yishak Woldetsadik conducted mouse work and antibody assays. Hsuan Liu, Harrison Powell, and Allison Chen created and titered the recombinant mouse-adapted H1N1 influenza viruses and developed the 3D images of HA trimers. Virus plaque assays and growth curves were performed by Kirsten Littlefield and Allison Chen. Santosh Dhakal, Robert Maul, Alice Mueller, and Han-Sol Park conducted all flow cytometry and data analyses. Zexu Ma was responsible for recombinant protein production and plasmid usage. Robert Maul conducted FACS sorting of germinal center B cells. Rebecca Ursin and Ashley Fink conducted B cell isolation and real-time RT-PCR. Sahana Jayaraman and H. Benjamin Larman performed the Phip-Seq. Han-Sol Park and Morgan Sherer conducted the immunofluorescence staining and imaging of mouse spleens. Rebecca Ursin, Santosh Dhakal, and Sabra Klein wrote the manuscript and all authors approved of the final draft.

## Acknowledgements

The authors thank Kumba Seddu, Aihui Wang, Georgia Stravrakis, and Maggie Li for assistance with assays and animal work. We thank Kim Davis for training and access to her laboratory’s cryostat and microscope. We are grateful to members of the Davis, Klein, and Pekosz labs for feedback on this work. Financial support provided by NIH/NIAID Johns Hopkins Center for Excellence in Influenza Research and Surveillance (JHCEIRS, contract # HHS N272201400007C, A.P. and S.L.K.) and the NIH/NIA Johns Hopkins Specialized Center of Research Excellence in sex and age differences in immunity to influenza (U54AG062333, S.L.K.). This work also was supported by NIH/NIAID T32A1007417 Molecular and Cellular Basis of Infectious Diseases (R.L.U.), NIH/NIGMS Hopkins PREP program R25GM109441 (Y.W.), and in part by the Intramural Research Program of the National Institute on Aging (R.W.M. and P.J.G.).

## Notes

### Competing Interest Statement

The authors have declared no competing interest.

