## Supplemental table and figure for "Greater breadth of vaccine-induced immunity in females than males is mediated by increased antibody diversity in germinal center B cells"

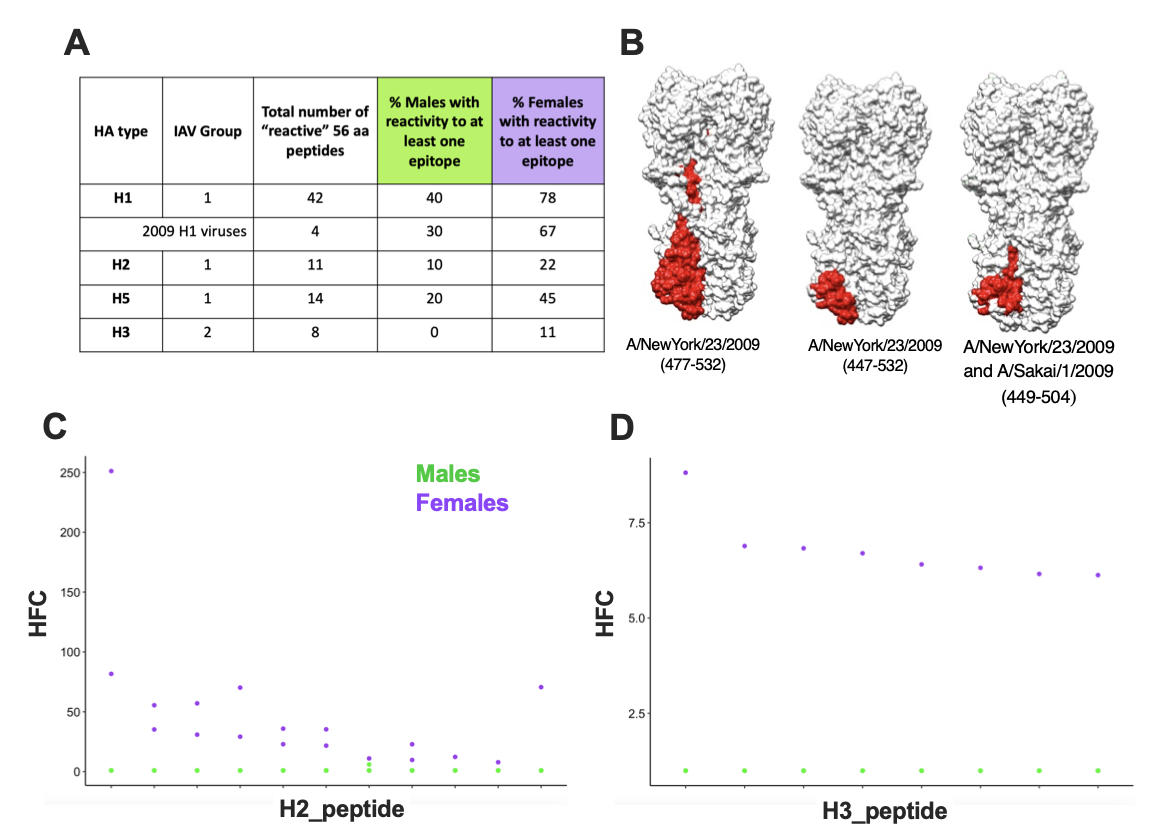

**Supplemental Figure 1: Vaccinated female mice exhibit greater reactivity to linear hemagglutinin stem-based epitopes as compared to male mice.** Serum from male and female mice (n=9-10/sex) vaccinated and boosted with mouse-adapted A/California/4/2009 (maA/Cal/09) H1N1 vaccine virus was used to determine antibody reactivity to thousands of linear influenza hemagglutinin (HA) epitopes via PhIP-Seq using the Vir-Scan library. A) The total number of reactive 56 amino acid (aa) peptides to unique H1, 2009-specific H1, H2 (group 1), H5 (group 1), and H3 (group 2) influenza A viruses (IAVs) having at least one animal with HFC > 1 are shown. Additionally, the percent of animals with at least one reactive epitope at 28 or 35 days post vaccination (i.e., 7 or 14 days post boost [dpb]) is indicated. B) The binding footprint of the four (three of which are unique) reactive A/Cal/09 H1N1 peptide amino acid sequences are shown on HA trimers and highlighted in red, with the corresponding virus name and amino acid positions shown below each structure. The A/Cal/09 protein was modeled as a trimer using the UCSF Chimera software and PDB model 3LZG. Reactivity of these vaccine-induced antibodies to unique H2 (C) and H3 (D) epitopes are shown, with each unique epitope represented on the X-axis and the hits fold-change (HFC) on the Y-axis. HFC is a measure of bound peptide in the serum-immunoprecipitated samples relative to the mock-immunoprecipitated samples, with no significant difference set to HFC = 1 (Methods).

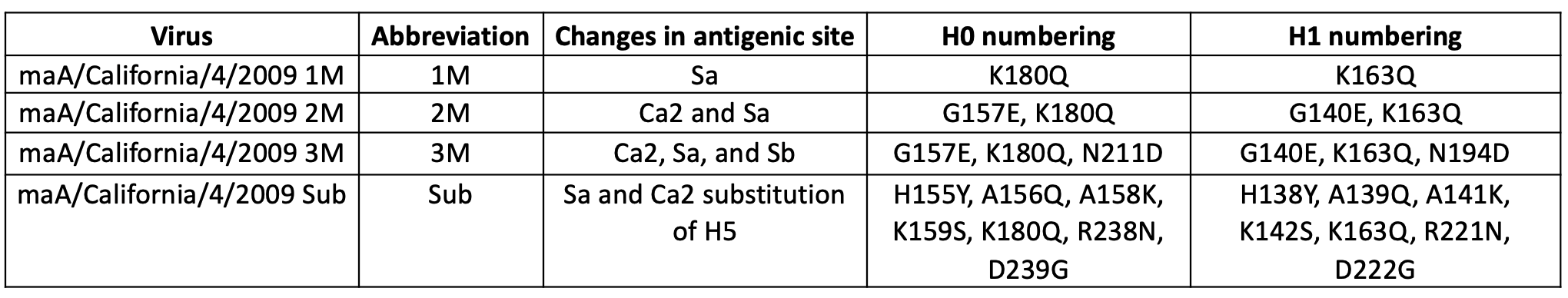

**Supplemental Table 1: Amino acid mutations on mouse-adapted A/California/4/2009 H1N1 virus hemagglutinin protein used in this study.** The parental mouse-adapted A/California/4/2009 (maA/Cal/09) H1N1 influenza virus was used to create four mutant viruses with either one, two, or three single point mutations in the hemagglutinin (HA) head (1M, 2M, 3M respectively), or a virus with an entire antigenic region substituted and replaced with a non-human H5 sequence (Sub). The antigenic sites affected by the mutations as well as the amino acids and numbering (for both H0 and H1) are listed for each mutant virus.
